# A Versatile *Agrobacterium*-Based Transformation System for Genetic Engineering of Diverse Citrus Cultivars

**DOI:** 10.1101/2022.06.30.498267

**Authors:** Michelle M. Dominguez, Carmen S. Padilla, Kranthi K. Mandadi

## Abstract

Developing an efficient transformation system is vital in genetically engineering recalcitrant crops, particularly trees. Here, we outline an *Agrobacterium tumefaciens*-based transformation methodology for citrus genetic engineering. The process was optimized to suit the requirements of fourteen citrus varieties by establishing appropriate infection, co-cultivation, selection, and culture media conditions. The procedure includes transforming seedling-derived epicotyl segments with an *A. tumefaciens* strain, then selecting and regenerating transformed tissues. Transgenic shoots were further identified by a visual reporter (e.g., β-glucuronidase) and confirmed by Northern and Southern blot analysis. Transgene integrations among the transgenic lines ranged between one to four. The methodology can yield transformation efficiencies of up to 11%, and transgenic plants can be recovered as early as six months, depending on the variety. In addition, we show that incorporating *A. tumefaciens* helper virulence genes (*vir*G and *vir*E), spermidine, and lipoic acid in the resuspension buffer before transformation improved the transformation efficiency of specific recalcitrant cultivars, presumably by enhancing T-DNA integration and alleviating oxidative stress on the explant tissues. In conclusion, the optimized methodology can be utilized to engineer diverse recalcitrant citrus varieties towards trait improvement or functional genetics applications.

## INTRODUCTION

Citrus is one of the most widely cultivated fruit crops in more than 130 countries in tropical and subtropical areas and is an important global commodity of significant economic value. Citrus and its products are rich in vitamins, minerals, and dietary fiber, essential for overall nutritional well-being. In 2019, citrus production was approximately 157.9 million tonnes worldwide, with oranges leading at 78.6 million, followed by tangerines, mandarins, clementines, satsumas (35.5 million), lemons and limes (20 million), and grapefruits (9.3 million) (FAO, 2019), but has been steadily declining due to unfavorable environmental conditions and diseases such as citrus canker and citrus greening (Albrecht et al., 2017).

The main objective of citrus breeding programs is to develop varieties with resistance or tolerance to problems such as pests, pathogens, drought, soil salinity, and many others. Conventional breeding has successfully improved citrus over the years. However, it has been met with obstacles due to the inherent biological limitations of citrus plants. Slow growth, long juvenile periods, regeneration time, nucellar polyembryony, high heterozygosity, and self-incompatibility make citrus one of the most difficult crops to breed. Many of these challenges with conventional breeding could be overcome through genetic engineering (Limera et al., 2017;Poles et al., 2020).

Genetic engineering in plants has been used for several decades (Fleming et al., 2000;Parisi et al., 2016). Currently, genetically modified crops in commercial production have had a positive global impact because of their resistance to pests and diseases, tolerance to pesticides, and exhibition of desirable nutritional traits. Examples of genetically modified crops include the glyphosate-resistant Roundup Ready^®^ Maize, soybean, and cotton from Monsanto (Funke et al., 2006;Lombardo et al., 2016;Parisi et al., 2016); virus-resistant papaya (Azad et al., 2014); virus-resistant plum (Fitch et al., 1992;Scorza et al., 2001;Scorza et al., 2007); non-browning apples (Waltz, 2015); and the golden rice with increased vitamin A content (Paine et al., 2005;Pérez-Massot et al., 2013).

In citrus, several methods for genetic transformation have been reported, which include particle bombardment (Yao et al., 1996), protoplast, shoot and root transformation using *Agrobacterium tumefaciens* or *Rhizobium rhizogenes* (Pena et al., 1995;Fleming et al., 2000;Yang et al., 2000;De Oliveira et al., 2008;Dutt et al., 2011;Yang et al., 2011;Sendin and Filippone, 2019;Irigoyen et al., 2020), however, *A. tumefaciens*-mediated transformation is the most widely used method. Because citrus is considered a recalcitrant species to genetic modification, the success of obtaining transgenic citrus largely depends on a reliable and empirically determined transformation and regeneration methodology. In the present work, we describe an optimized *A. tumefaciens*-based transformation protocol successfully used to generate transgenic plants from epicotyl tissue of fourteen different citrus varieties. We also highlight unique cultivar-specific transformation needs.

## MATERIALS AND METHODS

### Plant Material

Mature fruit was harvested from field-grown citrus trees and carefully cut open to manually extract the seeds. The seeds were washed with double deionized water until all the pulp and sugar were loosened and removed. Seeds were laid out to air dry at room temperature for up to 24 hours and then stored at 4°C. About 250 hydrated seeds were wrapped in three layers of damp paper towels and placed at 4°C overnight. Seeds were surface sterilized with 70% ethanol for two minutes and disinfected with two consecutive washes of 30% Clorox^®^, 0.2% Tween-20 solution, and 20% Clorox^®^, 0.2% Tween-20 solution for two and a half hours each. Seeds were rinsed three times with sterile double deionized water in 15-minute intervals. Under sterile conditions, the seeds’ outer coat and endosperm were removed, and the seeds were planted cut-side up in Petri dishes containing ½ MS media (½ MS, Table 1) (Figure 1A). Seeds were allowed to germinate at room temperature under dark conditions, then were carefully sub-cultured to 3×4 Magenta GA-7 vessels (Bio-World, Dublin, OH). The seeds remained under these conditions until epicotyls grew to approximately 7.5 cm in height, about four weeks, then transferred to a 16/8 h (light/dark) photoperiod for 14 days at 28°C.

**Table 1.** List of media used for *Agrobacterium-*based citrus transformation.

| Name of Medium | Abbreviation | Composition (1 L volume) |
| --- | --- | --- |
| ½ MS (Murashige and Skoog) Media | ½ MS | MS Basal Salts (2.15 g), sucrose (20.0 g), B5 vitamin (1 mL), pH 5.8, Gelrite (2.45 g) |
| ½ MS Liquid | ½ MSLIQ | MS Basal Salts (2.15 g), sucrose (25.0 g), MS vitamins (1 mL), pH 5.2, Lipoic Acid (5 µM)<sup>¶</sup>, Spermidine (1 mM)<sup>¶</sup> |
| DBA3 Co-Cultivation Media | CCM | MS Basal Salts (4.33 g), sucrose (25.0 g), malt extract (1.5 g), BAP* (1 – 3 mg), K<sub>2</sub>HPO<sub>4</sub> (20 mg), 2,4-D (10 ug), pH 5.8, Gelrite (2.5 g), coconut water (20 mL), B5 vitamin (1 mL), Acetosyringone (100 µM), Lipoic Acid (50 µM) |
| DBA3 Selection Media | SM | MS Basal Salts (4.33 g), sucrose (25.0 g), malt extract (1.5 g), BAP* (1 – 3 mg), K<sub>2</sub>HPO<sub>4</sub> (20 mg), 2,4-D (10 ug), pH 5.8, Gelrite (2.5 g), coconut water (20 mL), B5 vitamin (1 mL), Carbenicillin (200 mg), Cefotaxime (200 mg), Kanamycin (50 mg), Lipoic Acid (50 µM) |
| MS Shoot Elongation Media | SEM | MS Basal Salts (4.33 g), sucrose (25.0 g), B5 vitamin (1 mL), GA<sub>3</sub> (0.5 mg), BAP (0.2 mg), pH 5.8, Gelrite (2.0 g), Carbenicillin (100 mg), Cefotaxime (50 mg) |
| Rooting Media | RM | MS Basal Salts (2.15 g), B5 vitamin (1 mL), IAA (0.7 mg), pH 5.8, Gelrite (2.5 g) |
| E3P Charcoal Rooting Media | E3PRM | E3P Basal Salts w/ Vitamins and Adenine (3.98 g), sucrose (30.0 g), K-NAA* (2 – 5 mg), activated charcoal (1.0 g), pH 5.7, Gelrite (2.5 g) |
| DKW Regeneration Media<sup>#</sup> | DKW | DKW Basal Salts (5.22 g), sucrose (30.0 g), pH 5.7, Agar (8.0 g), BAP (1 mg) |
| MS Rooting Media<sup>#</sup> | MSRM | MS Basal Salts (4.33 g), sucrose (30.0 g), pH 5.7, Gelrite (2.5 g), IBA (2 mg), K-NAA (0.5 mg) |
| B5 Vitamin |  | Nicotinic Acid (0.1 g), Thiamine-HCl (0.1 g), Pyridoxine-HCl (0.1 g), Glycine (0.2 g), Myo-Inositol (10.0 g), double deionized sterile water to 100 mL volume. Filter sterilize. |

<sup>¶</sup> Included in the final Agro resuspension buffer. Spermidine was omitted for the sweet orange transformation
\*Dependent on the variety
<sup>#</sup>Carrizo Citrange only
BAP: 6-Benzylaminopurine; GA<sub>3</sub>: Gibberellic Acid; IAA: 3-Indoleacetic acid; K-NAA: Naphthalene acetic acid potassium salt; IBA: Indole-3 butyric acid.

**Figure 1.**
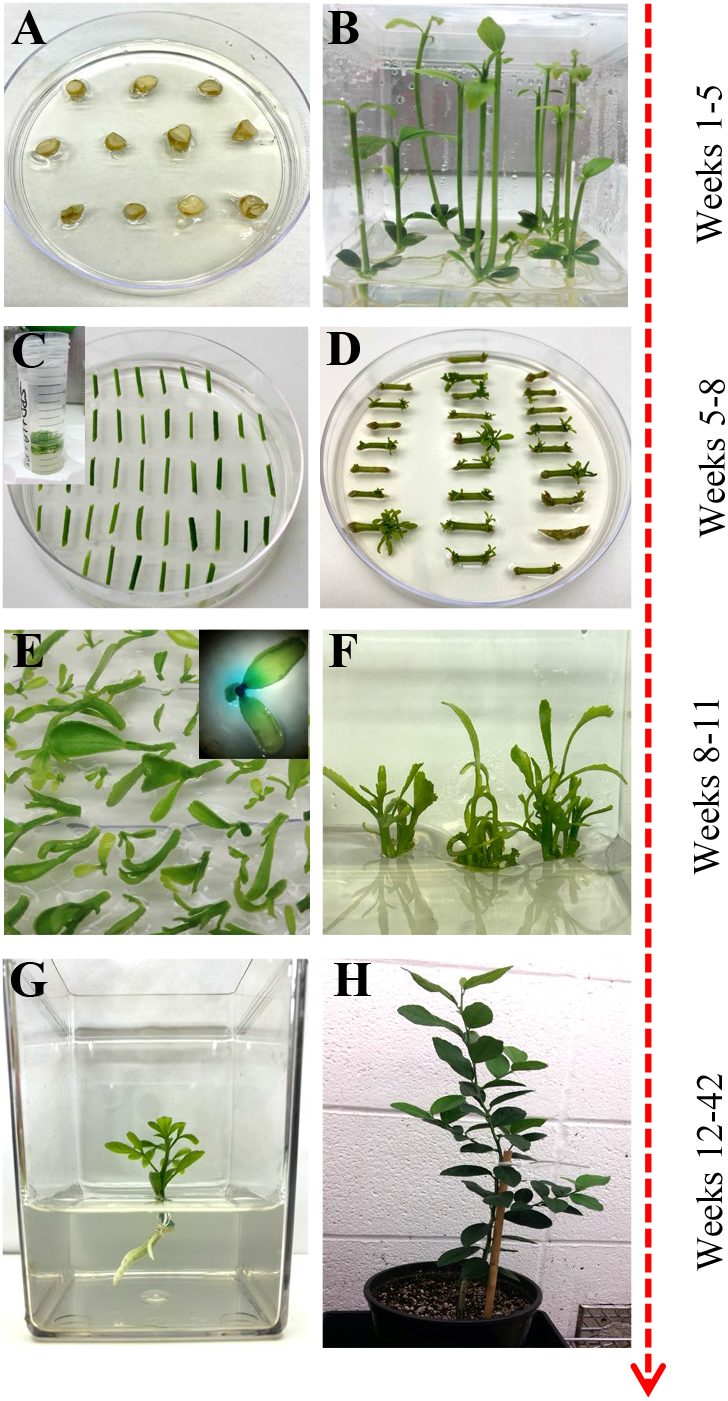
Overview of *Agrobacterium*-based citrus transformation. (**A-B**) Seed germination to epicotyl elongation; (**C**) Epicotyl *Agrobacterium-*based transformation and selection; (**D**) Shoot regeneration; (**E**) Reporter based screening (optional); (**F**) Shoot elongation and multiplication; (**G**) Root initiation; (**H**) Soil and greenhouse acclimation.

### Bacterial Culture

*A. tumefaciens* strain EHA105 (Hood et al., 1993) harboring pBIN34sGUS with the gene of interest (GOI) and a co-transformed plasmid (pCH32) containing *vir*G and *vir*E (Srivatanakul et al., 2000) was used for transformation (Supplementary Table 1). The stock was grown on YEP (yeast extract, peptone, sodium chloride) solid media supplemented with Tetracycline (5 mg/L), Kanamycin (50 mg/L), and Rifampicin (30 mg/L) one week before transformation at 28°C under dark conditions until single colonies formed (about three days). A single colony was selected and grown in 200 mL of liquid Luria Broth (LB) media supplemented with Tetracycline (5 mg/L), Kanamycin (50 mg/L), and Rifampicin (30 mg/L) at 28°C under dark conditions at 200 rpm one day before the transformation. Exponentially growing cells were centrifuged at 5,000 rpm for 20 minutes at 20°C. An optical density (OD_600_) of 0.6 to 1.0 was used for most varieties, except 0.3 was used for Mexican Lime (Supplementary Table 2). The cells were washed with ½ MS liquid media (Table 1) supplemented with 200 µM Acetosyringone, centrifuged again, and finally re-suspended in ½ MS liquid media. A final OD_600_ measurement was taken, and the culture was supplemented with 40 µM Acetosyringone and gently shaken at 160 rpm for 1 hour at 28°C under dark conditions. Lipoic acid (5 µM) and spermidine (1 mM) may be added to the bacterial cultures to enhance the transformation of most cultivars (Supplementary Tables 3 and 4)

### *A. tumefaciens*-mediated Transformation and Selection

Bright-green epicotyls were cut transversely into 1 cm length segments with angled, tapered ends to increase surface area and gently shaken in the *A. tumefaciens* suspension prepared above for 20 minutes at room temperature. The cell suspension was removed, and the explants were blotted dry on sterile filter paper to remove excess bacteria. Explants were placed horizontally on DBA3 co-cultivation media (CCM, Table 1) (Figure 1C) (Deng et al., 1992) and incubated at 22°C under dark conditions for three days. The explants were subsequently transferred to DBA3 selection media (SM, Table 1) and maintained for three weeks in a 16/8 h (light/dark) photoperiod at 28°C. Shoots began to develop after two weeks.

### Shoot Elongation and Reporter-based Screening

Regenerated shoots with green leaves approximately 4 mm in height were carefully excised from the explants and laid flat on Shoot Elongation Media (SEM, Table 1) (Yang et al., 2000) to be assayed for β-glucuronidase (GUS) activity if a GUS reporter gene was used in the constructs. Simultaneously, all healthy explants were sub-cultured onto SEM for further shoot regeneration and elongation. This process was repeated every three weeks for up to four months.

For GUS staining, a sterile X-Gluc solution (2 mM X-Gluc in 100 mM NaHPO_4_ Buffer, 0.1% Triton X-100) was added over the excised shoots and left at room temperature for up to 24 hours to stain. GUS-positive shoots were removed and carefully washed with double deionized sterile water and blotted dry on sterile filter paper. GUS-positive shoots were moved to SEM (Table 1) under a 16/8 h (light/dark) photoperiod at 28°C until the shoot grew approximately 1 to 2 cm in height.

### Rooting and Multiplication

The healthy GUS-positive shoots were top-grafted onto standard rootstock (Yang et al., 2000) or rooted. For root development, the callused base of the plantlet was sliced off before being placed in rooting media (RM, MSRM, Table 1) or charcoal supplemented rooting media (E3PRM, Table 1) (Albrecht et al., 2017) with its respective naphthalene acetic acid potassium salt (K-NAA) concentration (Table 2). The transgenic lines were ready for transplanting or *in vitro* micro-propagation when the plant was approximately 10 to 20 cm in height, around 6 to 8 weeks. For micro-propagation, briefly, the top half of the plant was decapitated, the leaves and thorns were removed, and the budwood was cut into nodal segments, with 2 mm and 6 mm of tissue above and below the node, respectively. The nodal segments were planted vertically into ½ MS media (Table 1) until buds began to develop, then sub-cultured to SEM (Table 1) for further shoot elongation (the callused base of the nodal segment was removed between each transfer). Carrizo Citrange nodal segments were planted and sub-cultured into DKW media (Table 1) (De Oliveira et al., 2016). Budded shoots were excised from the nodal segment when they were 1.5 cm tall and were rooted as described above.

**Table 2.**
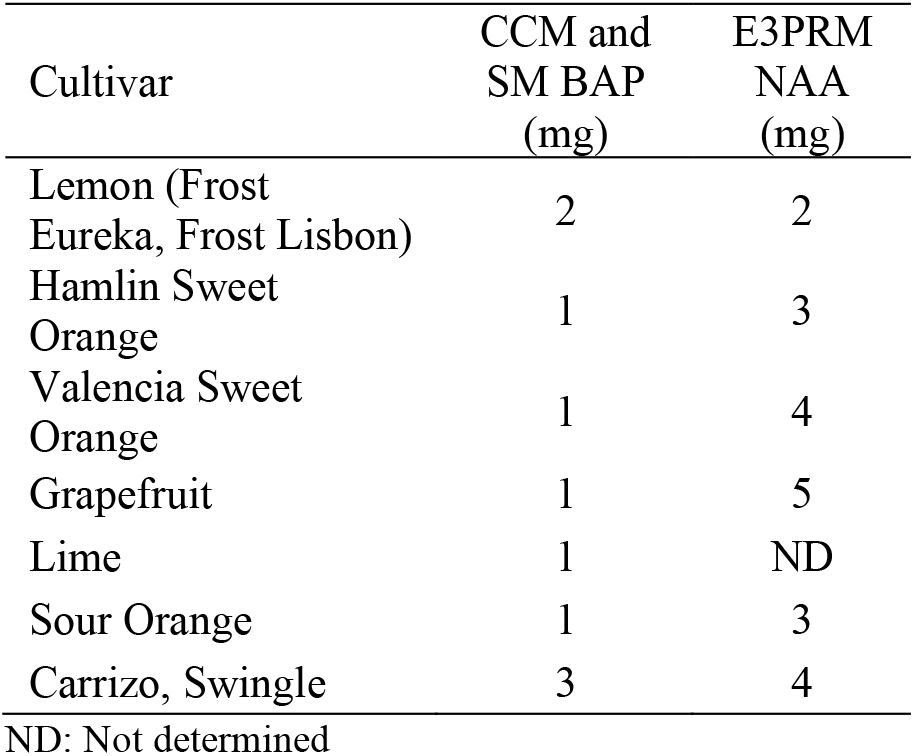
Final BAP and NAA concentrations used for co-cultivation, selection, and rooting media for the different varieties.

| Cultivar | CCM and<br>SM BAP<br>(mg) | E3PRM<br>NAA<br>(mg) |
| --- | --- | --- |
| Lemon (Frost<br>Eureka, Frost Lisbon) | 2 | 2 |
| Hamlin Sweet<br>Orange | 1 | 3 |
| Valencia Sweet<br>Orange | 1 | 4 |
| Grapefruit | 1 | 5 |
| Lime | 1 | ND |
| Sour Orange | 1 | 3 |
| Carrizo, Swingle | 3 | 4 |

ND: Not determined

### Potting and Acclimation

Plants were carefully removed from tissue culture, and the roots were washed with deionized water to remove any media. Plants were carefully planted into 1-pint pots that contained BG1 soil (Kinney Bonded, Donna, TX) and were loosely covered with moistened Ziploc bags to mimic a humid environment. The plants remained under a controlled environment (14/10 h light/dark photoperiod at 23°C) for approximately four weeks before being hardened under greenhouse conditions.

### Nucleic Acid Isolation

Total RNA was extracted from fresh leaf samples (1.5 g) of transgenic plants and control cultivars (Mangwende et al., 2009). Leaves were ground to a fine powder in liquid nitrogen, transferred to 50 mL conical tubes, and then frozen at −80°C. Three milliliters of HCl-Tris Saturated phenol were added and vortexed thoroughly before adding 6 mL of TENS Buffer (100 mM NaCl, 10 mM Tris-HCl, 1 mM EDTA, 1% SDS). The mixture was incubated for 5 minutes at 70°C with intermittent mixing every 2 minutes, then 3 mL of chloroform was added and vortexed for 1 minute. The samples were centrifuged at 10,000 rpm for 10 minutes, and 5 mL of the upper aqueous phase was carefully transferred to a new 50 mL tube. A 1:10 volume of 3 M DEPC-treated sodium acetate (pH 5.2) was added and gently mixed. Three volumes of cold 100% ethanol were added to the mixture, gently mixed, then incubated at −20°C for 2 hours. The samples were centrifuged at 10,000 rpm for 10 minutes at 4°C, the supernatant was discarded, and the pellet was resuspended with 5 mL DEPC-treated sterile double deionized water. Five milliliters of 4 M DEPC-treated lithium chloride were added, gently mixed, and left overnight at −20°C. The samples were centrifuged at 10,000 rpm for 20 minutes at 4°C, the supernatant was discarded, and the pellet was washed with cold 70% ethanol before being centrifuged again at 3,000 rpm for 5 minutes. The ethanol was removed, and the pellet was left to air dry before being dissolved in 200 µl DEPC-treated sterile double deionized water.

Genomic DNA was extracted from fresh leaf samples (0.8 g) of transgenic plants and control cultivars (Chee et al., 1991). Leaves were ground to a fine powder in liquid nitrogen, transferred to 15 mL conical tubes, and frozen at −80°C. Each sample was mixed with 8 mL of pre-warmed DNA Extraction Buffer (0.1 M Tris-HCl, pH 7.5; 0.7 M NaCl; 0.01 M EDTA, pH 8.0), 100 µl ß-mercaptoethanol, and 1.5 mL pre-warmed 10% CTAB. Samples were vortexed for 1 minute and incubated at 70°C for 90 minutes with intermittent mixing every 15 minutes. The mixture was allowed to cool for 5 minutes at room temperature before adding 4 mL of 24:1 chloroform: octanol. Samples were mixed by inversion for 5 minutes and then centrifuged at 5,000 rpm for 10 minutes at room temperature. Eight milliliters of the upper aqueous phase were transferred to a new 15 mL tube, and the chloroform:octanol step was repeated. After centrifugation, 7 mL of the upper aqueous phase was transferred to a 15 mL falcon tube, and an equal volume of cold isopropanol was added, gently mixed, and incubated for 30 minutes at − 20°C. Samples were centrifuged at 8,000 rpm for 5 minutes at room temperature, the supernatant was discarded, and the pellet was washed with 3 mL of 76% ethanol, 0.2 M sodium acetate for 20 minutes. The samples were centrifuged, the supernatant discarded, and washed with 1 mL of 76% ethanol, 10 mM ammonium acetate. After another centrifugation, the pellet was carefully transferred to a 1.5 mL tube using a glass hook and resuspended in 500 µl TE Buffer (50 mM Tris-Base, 10 mM EDTA, pH 8.0) and incubated with 10 µl RNAse A (10 mg/ mL) for 1 hour at 37°C. One hundred-fifty microliters of 5 M potassium acetate were added to each sample. Samples were shaken vigorously, incubated on ice for 20 minutes, and centrifuged at 14,000 rpm for 10 minutes at 4°C. The supernatant was transferred to a new 1.5 mL tube and incubated overnight at −20°C with 50 µl of 3 M sodium acetate (pH 5.2) and 550 µl of cold isopropanol. The samples were centrifuged at 8,000 rpm for 5 minutes, the supernatant discarded, and the pellet was washed with 1 mL of cold 70% ethanol. The samples were centrifuged again, ethanol discarded, and the pellet was dried at 37°C for 10 minutes before dissolving in 200 µl sterile double deionized water.

### Northern and Southern Blot Analysis

For Northern Blot analysis, ten micrograms of RNA were mixed with 7 µl of RNA and 2 µl ethidium bromide and incubated at 65°C for 4 minutes. The samples were placed on ice for 2 minutes, and 2 µl of bromophenol blue loading buffer was added. The samples were separated by gel electrophoresis on a 1.6% formaldehyde-agarose gel, then transferred to a positively charged nylon membrane by downward blotting in 20X SSC for 16 hours. The membrane was baked at 80°C for 15 minutes, UV crosslinked at 1,200 microjoules (x 100), and baked again at 80°C for 2 hours. The membrane was briefly washed with 5X SSC for 1 minute at room temperature, and the solution was discarded for hybridization.

For Southern Blot analysis, fifteen micrograms of DNA were digested overnight with *HindIII* for GUS and *AscI+PacI* for the GOI. After separation by gel electrophoresis for 21 hours on a 0.8% agarose gel, the gel was treated with 0.25 M HCl for 20 minutes, washed with 0.4 M NaOH for 20 minutes, and transferred to a positively charged nylon membrane by downward blotting in 0.4 M NaOH for 3 hours. The membrane was baked at 80°C for 20 minutes. The membrane was briefly washed with sterile double deionized water for 1 minute at room temperature, and the solution was discarded for hybridization.

Pre-hybridization, hybridization, washing, and detection were the same for Northern and Southern hybridization. The membrane was pre-hybridized with enough Church’s Buffer at 65°C for 4 to 16 hours. The GUS and GOI DNA probes were labeled with ^32^P-dCTP using a random primer labeling kit from Invitrogen (Waltham, MA). The mixture was incubated for 3 hours at room temperature, and unincorporated radioactive nucleotides were removed using a Sephadex G50 column. The labeled DNA probe was collected into a 1.5 mL tube, boiled for 5 minutes, added to the membrane in the Church’s buffer, and incubated for 16 hours at 65°C. The membrane was washed three times at 65°C as follows: 2X SSC, 0.5% SDS for 40 minutes; 1X SSC, 0.25% SDS for 25 minutes; 0.5X SSC, 0.125% SDS for 20 minutes. The membrane was covered in plastic wrap with excess wash buffer removed and then exposed to X-Ray film for 1 to 16 hours.

## RESULTS AND DISCUSSION

### Transformation overview and optimization of critical parameters

This study describes a detailed *A. tumefaciens* transformation method for fourteen citrus rootstock and scion varieties. As shown in Figure 1, seeds were germinated *in vitro* and allowed to grow for about five weeks, after which the epicotyls were cut transversely into 1 cm explants and exposed to the *A. tumefaciens*. All explants were subsequently moved into co-cultivation and selection media until shoots were about 0.5 cm tall and screened using a reporter gene (e.g., GUS). Positive shoots were maintained in shoot elongation and rooting media until they were ready for greenhouse acclimation. With this method, we were able to recover transgenic plants in ∼6 to 12 months, depending on the variety. Transformation efficiency varied between 0.2– 11.5%, as shown in Table 3.

**Table 3.** Transformation efficiency of different citrus varieties.

| Cultivar | Variety | Independent Experiment | per transformation |  | Transformation Efficiency Against Starting Material (%) | Transformation Efficiency Range | Approximate Duration to Recover Transgenic Plants in Pots | Average Copy Number |
| --- | --- | --- | --- | --- | --- | --- | --- | --- |
|  |  |  | Number of Epicotyl Segments | Number of Recovered Transgenic Shoots |  |  |  |  |
| Grapefruit | Rio Red | 1 | 183 | 21 | 11.5% | 3.2 – 11.5% | 11 months | 2.0 |
|  |  | 2 | 190 | 6 | 3.2% |  |  |  |
|  |  | 3 | 800 | 27 | 3.4% |  |  |  |
|  | Ruby Red | 1 | 531 | 2 | 0.4% | 0.4 – 3.1% | 10 months | 1.5 |
|  |  | 2 | 381 | 12 | 3.1% |  |  |  |
|  |  | 3 | 396 | 5 | 1.3% |  |  |  |
| Sweet Orange | Hamlin | 1 | 455 | 1 | 0.2% | 0.2 – 6.7% | 12 months | 1.6 |
|  |  | 2 | 30 | 2 | 6.7% |  |  |  |
|  |  | 3 | 113 | 3 | 2.7% |  |  |  |
|  | Valencia | 1 | 210 | 3 | 1.4% | 0.3 – 1.4% | 12 months | ND |
|  |  | 2 | 232 | 3 | 1.3% |  |  |  |
|  |  | 3 | 324 | 1 | 0.3% |  |  |  |
|  | Marrs | 1 | 360 | 5 | 1.4% | 0.8 – 1.4% | 10 months | 1.4 |
|  |  | 2 | 400 | 3 | 0.8% |  |  |  |
|  | Rohde Red | 1 | 110 | 3 | 2.7% | 1.1 – 2.7% | 10 months | 1.3 |
|  |  | 2 | 180 | 2 | 1.1% |  |  |  |
| Lemon | Frost Lisbon | 1 | 192 | 8 | 4.2% | 3.5 – 5.4% | 10 months | 1.9 |
|  |  | 2 | 831 | 45 | 5.4% |  |  |  |
|  |  | 3 | 684 | 24 | 3.5% |  |  |  |
|  | Frost Eureka Lemon | 1 | 716 | 35 | 4.9% | 2.2 – 4.9% | 10 months | 2.3 |
|  |  | 2 | 774 | 28 | 3.6% |  |  |  |
|  |  | 3 | 694 | 15 | 2.2% |  |  |  |
| Lime | Mexican Lime | 1 | 200 | 2 | 1.0% | 1.0 – 1.8% | 9 months | 1.0 |
|  |  | 2 | 600 | 9 | 1.5% |  |  |  |
|  |  | 3 | 712 | 13 | 1.8% |  |  |  |
| Rootstock | Carrizo Citrange | 1 | 325 | 26 | 8.0% | 5.4 – 8.0% | 6 months | 1.5 |
|  |  | 2 | 317 | 17 | 5.4% |  |  |  |
|  |  | 3 | 300 | 17 | 5.8% |  |  |  |
|  | Swingle | 1 | 250 | 17 | 6.8% | 5.1 – 6.8% | 6 months | 2.9 |
|  |  | 2 | 429 | 22 | 5.1% |  |  |  |
|  | Flying Dragon | 1 | 200 | 3 | 1.5% | 1.3 – 5.8% | 9 months | 3.5 |
|  |  | 2 | 150 | 2 | 1.3% |  |  |  |
|  |  | 3 | 275 | 16 | 5.8% |  |  |  |
|  | Sour Orange | 1 | 258 | 7 | 2.7% | 2.7 – 4.3% | 10 months | ND |
|  |  | 2 | 350 | 15 | 4.3% |  |  |  |
|  | C22 | 1 | 240 | 6 | 2.5% | 1.0 – 2.5% | 8 months | 1.1 |
|  |  | 2 | 210 | 2 | 1.0% |  |  |  |

ND: Not determined

We also optimized specific parameters to enhance the transformation, especially for recalcitrant varieties. It is well known that *A. tumefaciens* triggers basal plant immune responses (Zipfel et al., 2006;Tsuda et al., 2012;Raman et al., 2022), which can be a two-edged sword. It may reduce the transformation frequency of the infected cells and necrotize the explant tissues due to the production of excessive ROS and secondary metabolites. Different cultivars may have varying degrees of immune reactions to *A. tumefaciens*. As such, optimizing the *A. tumefaciens* optical density is needed to fine-tune the virulence and the plant immune responses to yield a sufficient number of transformed cells that can be regenerated into plants. The optical density (OD_600_) of *A. tumefaciens* optimal for transformation ranged from 0.3 to 1.0. We observed that while most of the cultivars responded well to an OD_600_ of 0.6 to 1.0, varieties such as Mexican Lime had better transformation efficiencies at the lower end of the range of 0.3 (Supplementary Table 2).

Next, to promote T-DNA integration and transformation, we utilized a helper plasmid (pCH32) containing *A. tumefaciens* virulence genes, *vir*G, and *vir*E (Srivatanakul et al., 2000) along with acetosyringone. In our experiments with Lisbon Lemon, we observed an approximately 50% increase in transgenic shoot recovery with the addition of the helper plasmid. The impact was similar to a previous report on its use in tobacco (Srivatanakul et al., 2000) and Clementine mandarin (Cervera et al., 2008). The number of transgenic shoots obtained with the pCH32 plasmid was 13 out of 327 explants (∼4%), and without the incorporation of pCH32 was 4 out of 236 explants (∼1.7%)(Supplementary Table 1).

Lipoic acid is an antioxidant, while spermidine is a polyamine that can alleviate plant dehydration stress. Spermidine can enhance the *A. tumefaciens* T-DNA integration (Khanna and Daggard, 2003;Kumar and Rajam, 2005;Dutt et al., 2011). Prior published citrus transformation studies have used Lipoic acid as an addition to the co-cultivation or selection media (Dutt et al., 2011). Still, there were no reports of spermidine use for citrus transformation. Hence, we tested whether adding lipoic acid or spermidine early in the transformation process, i.e., *A. tumefaciens* resuspension liquid media (½ MSLIQ) before transformation, could help improve the T-DNA integration and alleviate explant stress, especially for recalcitrant varieties. We first evaluated the effect of lipoic acid (5 µM) on the transformation of Rio Red and Flying Dragon. We observed a 3.5X improvement in transformation efficiency of Rio Red, and only a modest improvement in Flying Dragon (Supplemental Table 3). Next, we tested the effect of spermidine (1 mM) in the *A. tumefaciens* resuspension liquid media (½ MSLIQ) media for the transformation of Sweet orange, Swingle, Frost Lisbon, Frost Eureka, and Flying Dragon. We observed improvement in transformation efficiencies in most varieties, except in the case of sweet orange, where including spermidine had a negative impact (Supplemental Table 4). In summary, the combination of utilizing helper virulence genes, acetosyringone to prime *A. tumefaciens*, and lipoic acid and spermidine in the *A. tumefaciens* resuspension liquid media before explant transformation could be employed to promote transformation efficiency of multiple recalcitrant citrus varieties.

### Regeneration and growth media composition

After transformation, the growth media composition strongly influences the regeneration of transformed shoots. 6-Benzylaminopurine (BAP) and NAA in shoot regeneration and rooting media can impact transformation efficiency by promoting cell proliferation and differentiation. Due to variety-specific growth, and environmental and genetic variations, there may be variabilities in responses among different explant sources (Almeida et al., 2002;Poles et al., 2020). Several studies have employed varying concentrations between 1-5 mg/L of BAP or NAA to promote shoot and root regeneration in Hamlin, Valencia, Sour Orange, Rough Lemon, Mexican lime, and grapefruits (Bordón et al., 2000;Ghorbel et al., 2000b;Almeida et al., 2002;Mendes et al., 2008;Dutt et al., 2009;Rattanpal et al., 2011). Similarly, activated charcoal addition to rooting media can promote root development by adsorbing inhibitory substances, decrease phenolic oxidation, and can simulate soil conditions (Almudena et al., 2010). In our experiments, we utilized 1 mg/L BAP for Hamlin and Valencia sweet orange, Rio Red and Ruby Red grapefruit, and Mexican Lime and Sour Orange, while 2 mg/L for Frost Eureka and Lisbon, and 3 mg/L for Carrizo Citrange and Swingle Citrumelo rootstocks. For NAA, we have used it at 2 mg/L for Frost Eureka and Lisbon Lemon, 3 mg/L for Hamlin and Sour Orange, 4 mg/L for Valencia, Carrizo Citrange and Swingle Citrumelo, and 5 mg/L for Rio Red (Table 2). In summary, researchers must determine the optimal dosages of the various hormones and additives to promote shoot and root regeneration based on explant material sources. The dosage range of 1-5 mg/L for BAP and NAA can be used as a starting reference for optimization (Table 2).

### Molecular Characterization of Transgenic Lines

When a transformation is performed using plasmids containing a reporter gene such as GUS or green fluorescent protein (GFP), a visual screening could be performed to verify transgenic plants, as described in the materials and methods. Furthermore, selected transgenic lines can be evaluated using diagnostic tools such as Southern blot and Northern blot analysis to verify transgene integration and expression (Figure 2). Stable integration of transgene in the different citrus cultivars was determined by Southern blot hybridization. Average transgene integrations ranged from one to four integrations per transgenic line (Table 3, Fig. 2a). Figure 2b shows reporter gene expression levels and an example gene of interest (GOI). This was a heterologous plant defense gene that had no impact on the reported plant transformation or growth (Supplementary Figure 1) in four representative lines with a single transgene integration. Transgenic lines #1, 3, and 4 showed similar GUS expression levels after 1 hour of exposure, while transgenic line #2 showed lower expression (Fig. 2b, upper panel). GOI expression was similar between transgenic lines #3 and 4 after 3 hours of exposure (Fig. 2b, middle panel). To confirm the GOI expression of transgenic lines #1 and 2, a second exposure was performed for 16 hours (Fig. 2b, lower panel).

**Figure 2.**
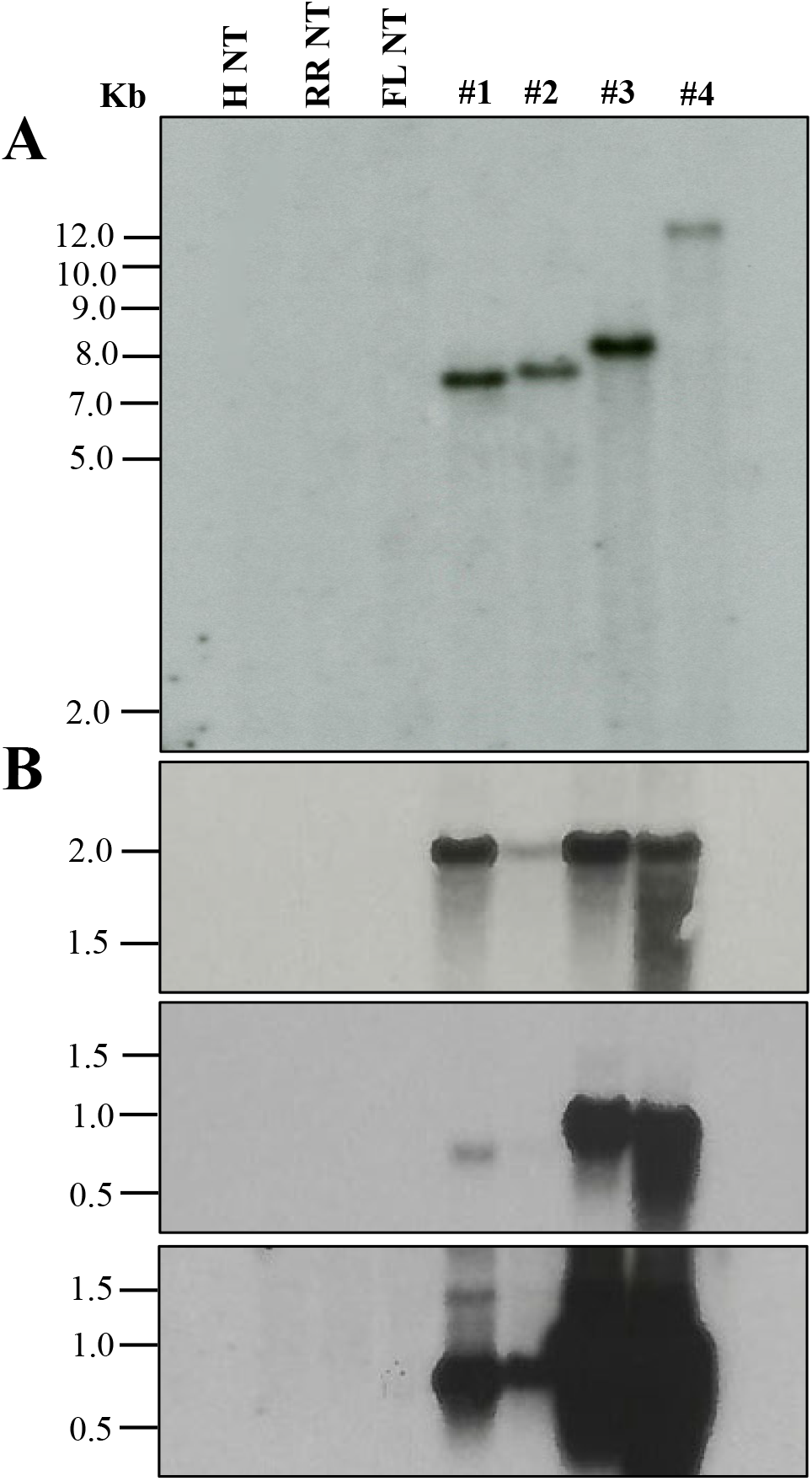
Molecular characterization of transgenic plants. (**A**) Southern blot analysis to assess transgene integration. (**B**) Northern blot analysis to evaluate transgene expression. GUS marker expression (upper panel), gene of interest (GOI) expression after 3 h exposure (middle panel) and after 16 h exposure (lower panel). Lanes: H NT-Hamlin non-transformed control, RR NT-Ruby Red non-transformed control, FL NT-Frost Lisbon non-transformed control, 1-Ruby Red transgenic, 2 and 3-Hamlin transgenic, 4-Frost Lisbon transgenic. Marker: 1 Kb Plus DNA Ladder (Invitrogen, Waltham, MA).

### Transformation Efficiency and Timeframe

Comparing transformation efficiencies (TE) and methods reported among different groups can be challenging. Critical points to consider are how the TE is defined, varietal differences, and the timeline. Regarding the TE definition, a common one is estimating the percentage of transgenic shoots recovered from starting epicotyl segments (Pena et al., 1995;Dutt and Grosser, 2009;Dutt et al., 2011). However, few studies also reported transformation frequencies, defined as the percentage of transformed events (at the cellular or calli level) in a given number of transformed explants (Pena et al., 1997;Ghorbel et al., 2000a). This distinction is essential to consider when comparing efficiencies because many cultivars can be successfully transformed at the cellular level; however, regenerating a transgenic shoot or plant can be difficult.

In this study, we report TE, defined as the percentage of recovered GUS-positive shoots/plants in a given number of transformed explants. Overall, among grapefruit varieties such as Rio Red and Ruby Red, the transformation efficiency was between 3.2–11.5% and 0.4– 3.1%, respectively. Among the sweet orange varieties, Hamlin exhibited a better efficiency range of 0.2–6.7%, while transformations with Valencia, Marrs, and Rohde Red showed 0.3–1.4%, 0.8–1.4%, and 1.1–2.7%, respectively. In the lemons and lime tested, Frost Lisbon transformation efficiency ranged from 3.5–5.4%, Frost Eureka Lemon ranged from 2.2–4.9%, and Mexican Lime ranged from 1.0–1.8%. Carrizo Citrange had the highest range of transformation efficiency among the rootstocks evaluated at 5.4–8.0%, followed by Swingle at 5.1–6.8%, Flying Dragon at 1.3–5.8%, Sour Orange at 2.7–4.3%, and C22 at 1.0–2.5%.

Regarding the timeline, studies have reported up to 6 months to recover transgenic citrus shoots, primarily influenced by the transformed variety and methodology employed (Pena et al., 1995;Fleming et al., 2000;Yang et al., 2000;De Oliveira et al., 2008;Dutt et al., 2011;Yang et al., 2011;Sendin and Filippone, 2019). For instance, Pena et al. (1995) reported screening transformed carrizo citrange shoots at ∼12 weeks after transformation. Two GUS-positive shoots out of 368 transformed explants were recovered (∼0.5 % TE). However, continued selection and screening for up to ∼6 months yielded up to ∼20% TE (Pena et al., 1995). Using the current method, we recovered an average of ∼20 GUS-positive shoots out of 300 transformed explants (∼6.6% TE, Table 3) in as little as 6 weeks. One could potentially recover more transgenic shoots if we continue screening explants/shoots for longer period, especially with Carrizo, which is prolific in regeneration. Ultimately, researchers must decide when to stop screening based on their experimental needs and timeframe.

## Conclusion

The current methodology was demonstrated for successful genetic engineering of a broad range of citrus varieties (14 varieties), including several recalcitrant varieties previously not transformed. We also note our observations and improvements that could be used to promote transformation efficiency. These include utilizing helper virulence genes to prime *A. tumefaciens* and adding spermidine and lipoic acid early on in transformation to the *A. tumefaciens* resuspension liquid media to enhance T-DNA integration and alleviate explant stress. The methodology can be a helpful guide for potential users working on one of these fourteen or other cultivars.

## Acknowledgments

The authors would like to thank Denise Rossi, Sonia del Rio, Joe Molina, Rolando Mireles Kinnie Laughlin, Manikandan Ramasamy, Shreya Udawant (Texas A&M AgriLife Research), Mike Irey (Southern Gardens Citrus, FL), Beth Lamb (Philip Rucks Nursery, FL) and TAMU-Kingsville citrus center for providing various citrus resources, technical assistance, and critical comments during manuscript preparation. The authors would like to dedicate this manuscript to Prof. T. Erik Mirkov (1959-2018), a renowned scientist in plant biotechnology. This study was supported in part by funds from USDA-NIFA (2021-70029-36056; HATCH TEX09621), Texas A&M AgriLife Research Insect-vectored Disease Seed Grants (124185-96210), the Texas A&M AgriLife Institute for Advancing Health Through Agriculture, and Southern Gardens Citrus (FL) to KM.

## Author contributions

MD and CP performed the experiments and analyzed the data. KM supervised the experiments, and all authors contributed to the preparation of the manuscript.

## FIGURE LEGENDS

**Supplementary Figure 1.**
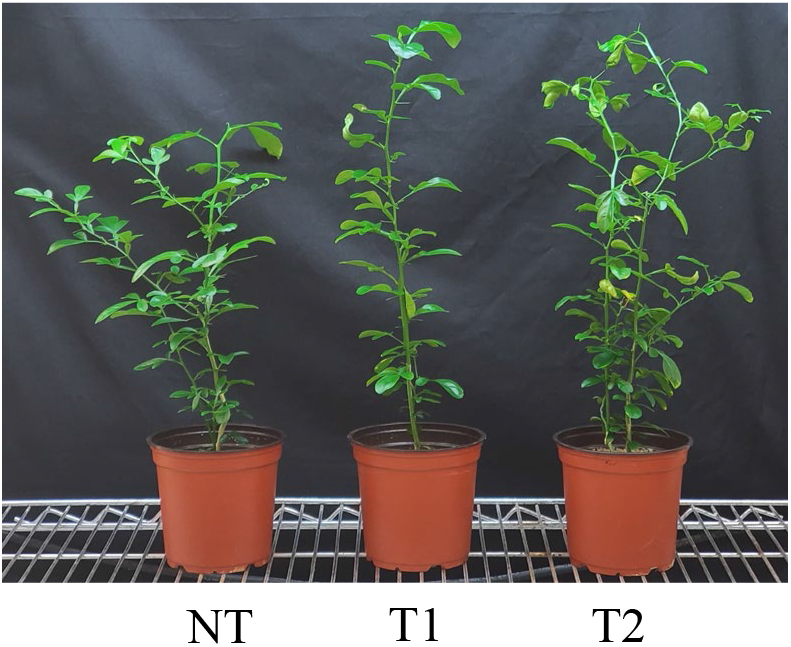
Comparison of one-year-old Carrizo Citrange non-transgenic (NT) and two independent transgenic lines with a heterologous gene of interest (T1 and T2).

**Supplementary Table 1.** Effects of pCH32 on transformation efficiency.

| Variety | Number of<br>GUS Positive<br>Shoots | Total<br>Number of<br>Explants | Transformation<br>Efficiency<br>(%) |
| --- | --- | --- | --- |
| Frost Lisbon – pCH32 | 4 | 236 | 1.7 |
| Frost Lisbon + pCH32 | 13 | 327 | 4.0 |

**Supplementary Table 2.** Effects of *A. tumefaciens* density (OD_600_) on transformation efficiency.

| Variety | OD<sub>600</sub> | Number of<br>GUS<br>Positive<br>Shoots | Total<br>Number of<br>Explants | Transformation<br>Efficiency<br>(%) |
| --- | --- | --- | --- | --- |
| Frost Lisbon | 0.3 | 4 | 209 | 1.9 |
|  | 0.6 | 8 | 204 | 3.9 |
|  | 1.0 | 7 | 230 | 3.0 |
| Sour Orange | 0.3 | 8 | 420 | 1.9 |
|  | 0.6 | 6 | 264 | 2.3 |
|  | 1.0 | 0 | 502 | 0.0 |
| Mexican Lime | 0.3 | 22 | 1312 | 1.7 |
|  | 0.6 | 0 | 262 | 0.0 |
|  | 1 | 0 | 600 | 0.0 |

**Supplementary Table 3.** Effects of Lipoic Acid (LA) on transformation efficiency.

| Variety | Number of<br>GUS Positive<br>Shoots | Total Number<br>of Explants | Transformation<br>Efficiency<br>(%) |
| --- | --- | --- | --- |
| Rio Red – LA | 6 | 495 | 1.2 |
| Rio Red + LA | 21 | 495 | 4.2 |
| Flying Dragon – LA | 3 | 300 | 1.0 |
| Flying Dragon + LA | 4 | 300 | 1.3 |

**Supplementary Table 4.** Effects of Spermidine (SPD) on transformation efficiency.

| **Cultivar** | **Number of**<br>**GUS Positive**<br>**Shoots** | **Total Number**<br>**of Explants** | **Transformation**<br>**Efficiency**<br>**(%)** |
| --- | --- | --- | --- |
| Swingle – SPD | 84 | 1300 | 6.5 |
| Swingle + SPD | 73 | 1404 | 5.2 |
| Flying Dragon – SPD | 12 | 360 | 3.3 |
| Flying Dragon + SPD | 22 | 384 | 5.7 |
| Frost Lisbon – SPD | 12 | 480 | 2.5 |
| Frost Lisbon + SPD | 21 | 525 | 4.0 |
| Frost Eureka – SPD | 3 | 300 | 1.0 |
| Frost Eureka + SPD | 27 | 561 | 4.8 |
| Sweet orange – SPD | 21 | 748 | 2.8 |
| Sweet orange + SPD | 0 | 1594 | 0.0 |

